# Individualized isometric neuromuscular electrical stimulation training promotes myonuclear accretion in mouse skeletal muscle

**DOI:** 10.1101/2021.12.14.472254

**Authors:** Aliki Zavoriti, Aurélie Fessard, Masoud Rahmati, Peggy Del Carmine, Bénédicte Chazaud, Julien Gondin

**Affiliations:** Institut NeuroMyoGène, CNRS UMR 5310, INSERM U1217, Université Claude Bernard Lyon 1, Univ Lyon, Lyon, France; Department of Exercise Physiology, Faculty of Literature and Human Sciences, Lorestan University, Khoramabad, Iran

**Author notes:** Corresponding author: Julien GONDIN, Institut NeuroMyoGène (INMG), CNRS 5310 – INSERM U1217 – UCBL1, Faculté de Médecine et de Pharmacie, 8 Avenue Rockefeller, 69008 LYON, FRANCE.

**Keywords:** satellite cells, force, *in vivo*, standardized training protocol

## Abstract

Skeletal muscle is a plastic tissue that adapts to exercise through fusion of muscle stem cells (MuSCs) with myofibers, a physiological process referred to as myonuclear accretion. However, it is still unclear whether myonuclear accretion is driven by increased mechanical loading *per se*, or occurs, at least in part, in response to exercise-induced muscle injury. Here, we developed a carefully monitored and individualized neuromuscular electrical stimulation (NMES) training protocol of the mouse plantar flexor muscles. Each NMES training session consisted of 80 isometric contractions at a submaximal mechanical intensity corresponding to ~ 15% of maximal tetanic force to avoid muscle damage. NMES trained mice were stimulated for 2 × 3 consecutive days separated by one day of rest, for a total of 6 sessions. Experiments were conducted on C57BL/6J and BALB/c males at 10-12 weeks of age. NMES led to a robust myonuclear accretion and higher MuSC content in *gastrocnemius* muscle of both mouse lines, without overt signs of muscle damage/regeneration or muscle hypertrophy or force improvement. This new mouse model of myonuclear accretion relying on the main function of skeletal muscles, *i.e.*, force production in response to electrical stimuli, will be of utmost interest to further understand the role of MuSCs in skeletal muscle adaptations.

## Introduction

Skeletal muscle is a remarkably plastic tissue that both regenerates *ad integrum* after an acute injury, and adapts to changes in mechanical loading (*e.g.*, disuse, overloading). This plasticity widely relies on muscle stem cells (aka satellite cells, MuSCs) which are located beneath the basal laminal, *i.e.*, on the periphery of the myofibers. While MuSCs are indispensable for muscle regeneration (Lepper *et al*, 2011; Murphy *et al*, 2011; Sambasivan *et al*, 2011), during which they exit quiescence, expand, differentiate and fuse to form new functional myofibers, emerging evidence also illustrates their roles in skeletal muscle homeostasis and remodeling (Murach *et al*, 2021).

Over the last few years, the contribution of MuSCs to the homeostasis of healthy myofibers has been reported, as illustrated by their fusion to uninjured myofibers in an age-, muscle- and myofiber-type-dependent manner (Keefe *et al*, 2015; Pawlikowski *et al*, 2015). Thanks to the recent development of genetic mouse models either ablated for MuSCs (Egner *et al*, 2016; McCarthy *et al*, 2011) or deleted for *myomaker* that is necessary for fusion (Goh & Millay, 2017), the role of MuSC fusion-induced myonuclear accretion has been further investigated in the context of increased mechanical loading. The requirement of MuSC-mediated myonuclear accretion for hypertrophy was demonstrated in young animals (< 4 months) (Egner *et al*, 2016; McCarthy *et al*, 2011; Goh & Millay, 2017) but not in mature animals (Murach *et al*, 2017) in a drastic model of overload relying on surgical ablation of synergist muscles. However, the physiological relevance of this model has been questioned not only regarding the magnitude of hypertrophy, that largely exceeds what can be achieved in humans after resistance training, but also due to the confounding effects of overload-induced muscle damage on the regulation of MuSC fate (Murach *et al*, 2017; Fukuda *et al*, 2019; Egner *et al*, 2016). To counteract these limitations, weighted voluntary wheel running (Dungan *et al*, 2019; Masschelein *et al*, 2020), high-intensity interval treadmill (Goh *et al*, 2019) or weight pulling (Zhu *et al*, 2021) protocols have been recently introduced to decipher the contribution of MuSCs to exercise-induced myonuclear accretion. Myonuclear accretion is dependent on the training load (Masschelein *et al*, 2020) and occurs early during training (Goh *et al*, 2019; Englund *et al*, 2021) while MuSC depletion blunts myofiber hypertrophy (Englund *et al*, 2021). Although these physiological models of exercise greatly contributed to improve our understanding on the role of MuSCs in skeletal muscle adaptations, unaccustomed running activity is known to induce muscle damage (Wernig *et al*, 1990; Irintchev & Wernig, 1987) due to the eccentric muscle contraction component of running. In addition, exercise design usually involves the same absolute increment of wheel load (Dungan *et al*, 2019) or running speed (Goh *et al*, 2019) for all mice so that the training load is not adjusted according to the individual performance capacity. The lack of running exercise individualization might further aggravate the extent of muscle damage (Murach *et al*, 2020a; Goh *et al*, 2019). As a consequence, myonuclear accretion might not only be driven by increased mechanical loading in these models, but may occur, at least in part, in response to running exercise-induced muscle injury.

Here, we developed a new and individualized resistance training protocol which relies on the main function of skeletal muscle, *i.e.*, its ability to produce force in response to repeated electrical stimuli. Thanks to our large experience in neuromuscular electrical stimulation (NMES) training (Gondin *et al*, 2005, 2006, 2011b), we designed a short NMES training protocol that was performed under isometric conditions to avoid muscle damage (Gondin *et al*, 2011a). In addition, the force produced by the plantar flexor muscles was adjusted to reach a submaximal level (*i.e.*, around 15% of maximal force) and was carefully monitored in response to each stimulation train and for each trained mouse. This innovative training modality led to a robust myonuclear accretion and higher MuSC content in two mouse lines with a different genetic background, without overt signs of muscle damage/regeneration. This new mouse model of myonuclear accretion will be of interest to further understand the role of MuSCs in skeletal muscle adaptations.

## Materials and Methods

### Animals

Experiments were conducted on both C57BL/6J and BALB/c males (Janvier Labs, Le Genest-Saint-Isle, France) at 10-12 weeks of age. These two genetic backgrounds are commonly used to investigate the impact of cancer cachexia (Gallot *et al*, 2014; Penna *et al*, 2016) or sepsis (Morel *et al*, 2017) on skeletal muscle homeostasis. Mice were housed in an environment-controlled facility (12-12 hour light-dark cycle, 25°C), received water and standard food *ad libitum*. All of the experiments and procedures were conducted in accordance with the guidelines of the local animal ethics committee of the University Claude Bernard Lyon 1 and in accordance with French and European legislation on animal experimentation and approved by the ethics committee CEEA-55 and the French ministry of research (APAFIS#12794-2017122107228405).

### Experimental device

In order to propose individualized and carefully monitored NMES training protocols, we used a strictly non-invasive ergometer (NIMPHEA_Research, AII Biomedical SAS, Grenoble, France) offering the possibility to electrically stimulate the plantar flexor mouse muscles and to record the resulting force production (Fig. 1A). Mice were initially anesthetized in an induction chamber using 4% isoflurane. The right hindlimb was shaved before an electrode cream was applied over the plantar flexor muscles to optimize electrical stimulation. Each anesthetized mouse was placed supine in a cradle allowing for a strict standardization of the animal positioning in ~1 min (Supplemental Video 1). Throughout a typical experiment, anesthesia was maintained by air inhalation through a facemask continuously supplied with 1.5-2.5% isoflurane. The cradle also includes an electrical heating blanket in order to maintain the animal at a physiological temperature during anesthesia. Electrical stimuli were delivered through two electrodes located below the knee and the Achille’s tendon. The right foot was positioned and firmly immobilized through a rigid slipper on a pedal of an ergometer allowing for the measurement of the force produced by the plantar flexor muscles (*i.e.*, mainly the *gastrocnemius* muscle). The right knee was also firmly maintained using a rigid fixation in order to optimize isometric force recordings.

**Figure 1.**
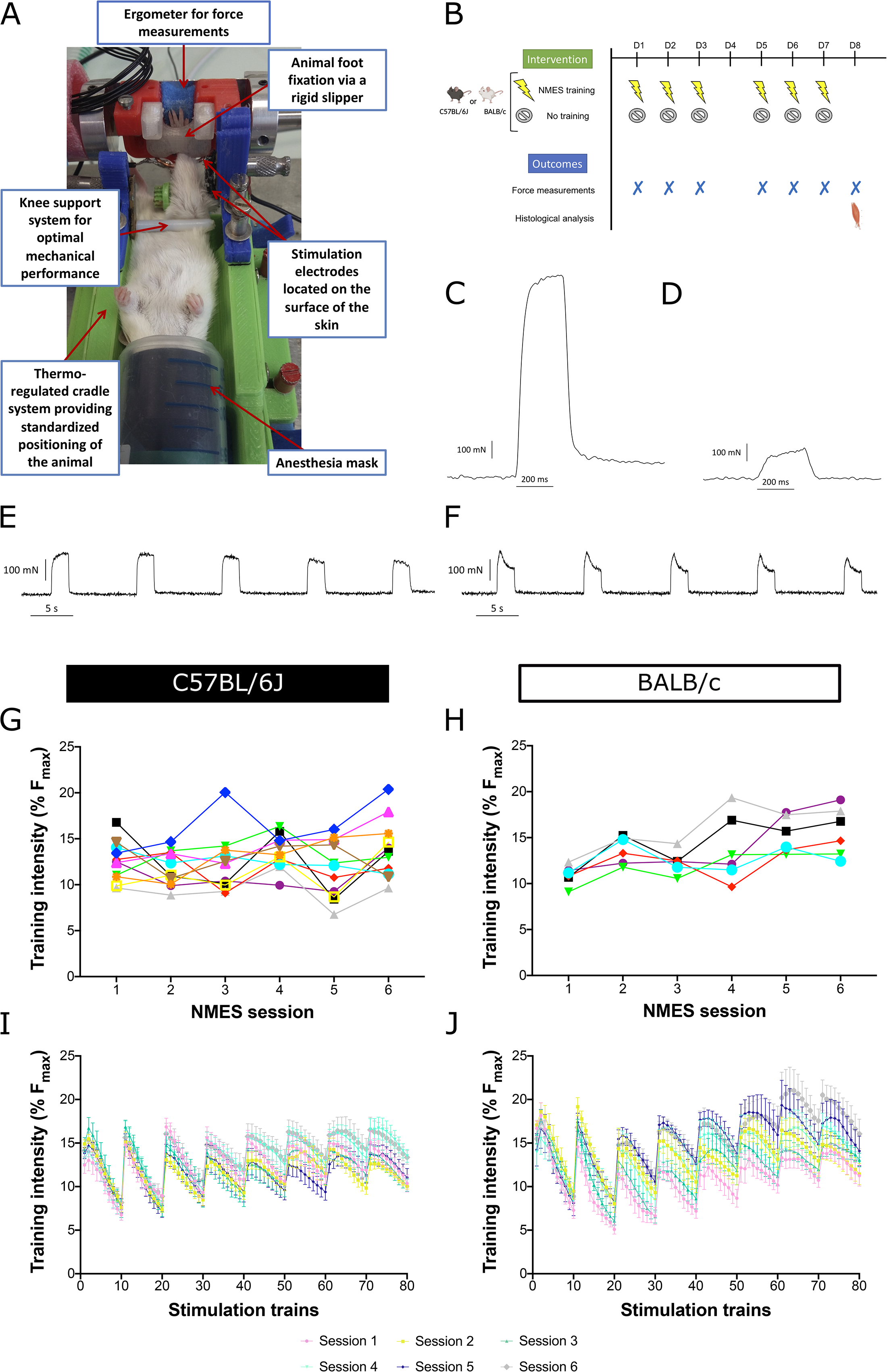
**A)** Experimental device allowing for non-invasive longitudinal force measurements and individualized neuromuscular electrical stimulation (NMES) training protocol in response to electrical stimuli applied over the plantar flexor muscle belly. **B)** Schematic representation of the study design. C57BL/6J or BALB/c were submitted to either an individualized NMES training protocol or a control intervention over an 8-day period. Maximal force production was recorded at the beginning of each NMES or control interventions as well as at day 8. Then, animals were sacrificed and *gastrocnemius* muscle was harvested. **C)** Typical mechanical trace obtained in response to a 250-ms 100 Hz tetanic stimulation train allowing for maximal isometric force production measurement. **D)** Typical mechanical trace obtained in response to a 250-ms 50 Hz subtetanic stimulation train allowing for the determination of the training intensity corresponding to 15% of maximal isometric force. **E-F)** Typical mechanical traces obtained during the first five (panel E) and last five (panel F) stimulation trains during a typical NMES training session. **G-H)** Individual training intensity (expressed in percentage of maximal isometric force) obtained during each NMES training session in C57BL/6J (n=11, panel G) and BALB/c mice (n=6, panel H). **I-J)** Mean training intensity (expressed in percentage of maximal isometric force) obtained during each NMES training session throughout the 80 stimulation trains in C57BL/6J (n=11, panel I) and BALB/c mice (n=6, panel J).

### *In vivo* maximal force measurements and NMES training

C57BL/6J and BALB/c males were submitted either to a NMES training protocol (*i.e.*, NMES mice) or a control intervention (*i.e.*, control mice) (Fig. 1B).

In both NMES and control mice, transcutaneous stimulation was first elicited on the plantar flexor muscles using a constant-current stimulator (Digitimer DS7AH, Hertfordshire, UK; maximal voltage: 400 V; 0.2 ms duration, monophasic rectangular pulses). The individual maximal current intensity was determined by progressively increasing the current intensity until there was no further peak twitch force increase. This intensity was then maintained to measure maximal isometric force production (F_max_) in response to a 250-ms 100 Hz tetanic stimulation train (Fig. 1C).

NMES mice were then submitted to a NMES protocol performed under isometric conditions at a submaximal mechanical intensity corresponding to ~15% of F_max_ in order to i) avoid muscle damage (Gondin *et al*, 2011a); ii) mimic the application of NMES in severely impaired patients (Maddocks *et al*, 2016) for whom higher force levels are difficult to reach due to discomfort associated with electrical stimuli (Gondin *et al*, 2011c). As a consequence, the current intensity was carefully adjusted at the beginning of each NMES training session in order to reach 15% of F_max_ (*i.e.*, initial intensity I_15%_; range: 12.5-17.5% of F_max_) in response to a 250-ms 50 Hz stimulation train (Fig. 1D). Each NMES session consisted of 80 stimulation trains (2-s duration, 8-s recovery) delivered at a frequency of 50 Hz. Every 10 contractions, the current intensity was increased by 50% from I_15%_ in order to minimize muscle fatigue and maintain a force level of ~15% of F_max_ throughout the NMES protocol (Fig. 1E-F). Current intensity (in mA) was consistently recorded and averaged for all stimulation trains for each training session. NMES mice were stimulated for 2 × 3 consecutive days separated by one day of rest, for a total of 6 sessions (Fig. 1B) corresponding to a total muscle contractile activity of only 16 min (*i.e.*, 6 sessions x 80 trains x 2 sec = 960 s). For all NMES training sessions and for each mouse, the force produced in response to each stimulation train was quantified and normalized to F_max_ recorded at the beginning of the corresponding session. Control mice were not stimulated but were kept under anesthesia for the same duration as an NMES session. At day 8, F_max_ was recorded in both NMES and control mice (Fig. 1B). Force signal was sampled at 1000 Hz using a Powerlab system and Labchart software (ADinstruments).

### Tissue preparation and immunofluorescence analyses

At day 8, all animals were sacrificed by cervical dislocation after deep isoflurane anesthesia. The right *gastrocnemius* muscle was harvested, weighted and then frozen in isopentane placed in liquid-nitrogen, and kept at −80°C until use. Cryosections (10 μm) were prepared for immunohistochemical analyses.

Cryosections were permeabilized in Triton-X100 0.5% for 10 min at room temperature, washed 3 times in PBS and then blocked in BSA 4% for 1 hour at room temperature. Cryosections were then incubated with primary antibodies overnight at 4°C, washed 3 times with PBS and further incubated with secondary antibody for 1 hour at 37°C. The following primary antibodies were used: anti-MYH3 (1/200, mouse, sc-53091, Santa Cruz Biotech) and anti-Laminin (1/200, rabbit, L9393, Merck). Secondary antibodies were: Alexa Fluor 488 AffinePure Goat Anti-Mouse (1/200, ref: 115-545-205), Cy3 AffinePure Donkey Anti-Rabbit (1/200, ref: 711-165-152), Cy3 AffinePure Donkey Anti-Mouse (*i.e.*, for determining IgG^+^ myofibers; 1/200, ref: 715-165-150), Fluorescein (FITC) AffinePure Donkey Anti-Rabbit (1/200, ref: 711-095-152) supplied from Jackson ImmunoResearch. Slides were washed with PBS, counterstained with Hoechst and mounted in Fluoromount-G medium

For Pax7 immunostaining, cryosections were first fixed with PFA 4% for 10 min, washed 3 times in PBS, permeabilized in Triton-X100 0.1% + 0.1M Glycine for 10 min, washed 3 times in PBS, then immerged into citrate buffer 10mM in 90°C hot water bath twice, washed 3 times in PBS and blocked in donkey serum 5% BSA 2% and MOM 1/40 for 1 hour. Every step was performed at room temperature, unless indicated otherwise. Cryosections were then incubated with antibodies as described above except that primary (anti-Pax7; 1/50, mouse, DSHB) and secondary antibodies were diluted in blocking buffer containing donkey serum 5% and BSA 2%.

### Image capture and analysis

Ten to fifteen images were recorded from each section with an Imager Z1 Zeiss microscope at 20x magnification connected to a CoolSNAP MYO camera for the quantification of the number of myonuclei per fiber and Pax7^+^ cells. Nuclei with their geometric center within the inner rim of the laminin ring were defined as myonuclei. The number of myonuclei was divided by the number of fibers analyzed on the same picture (Egner *et al*, 2016). The number of Pax7^+^ cells was also divided by the number of fibers analyzed on the same picture (Theret *et al*, 2017). For whole cryosection analysis, slides were automatically scanned at × 10 of magnification using an Axio Observer.Z1 (Zeiss) connected to a CoolSNAP HQ2 CCD Camera (photometrics). The image of the whole cryosection was automatically reconstituted in MetaMorph Software (Desgeorges *et al*, 2019).

The number of IgG positive fibers (*i.e.*, based on staining with cy3 anti-mouse), embryonic MyHC positive fibers and myofibers with central nuclei were quantified on the whole section and normalized to the total number of myofibers. Myofiber cross-sectional area (CSA) was determined on whole gastrocnemius muscle sections labeled by anti-laminin antibody using the Open-CSAM program, as previously described (Desgeorges *et al*, 2019).

### Statistical Analysis

Statistical analysis was performed using GraphPad Prism Software (version 9.0). Data distribution was initially investigated using Kolmogorov-Smirnov test. Two-factor (group x time) analysis of variance (ANOVAs) with repeated measures on time was used to compare maximal tetanic force. One-factor ANOVA with repeated measures on session was used to compare training intensity and current intensity. Unpaired student t-test was used to test differences between control and NMES mice for other variables. Data are presented as mean ± SD with significance set at p < 0.05

## RESULTS

### Individualization of NMES training program

Thanks to our original device allowing for longitudinal force recordings in response to electrical stimulation applied on the surface of the plantar flexor muscles (Fig. 1A & Supplemental video 1), F_max_ was recorded at the beginning of each NMES training session (Fig. 1B) for each NMES trained mouse (Fig. 1C). Then, the current intensity was carefully adjusted to reach 15% of F_max_ on the basis of a 250-ms testing train delivered at 50 Hz (Fig. 1D). The corresponding current intensity was applied for the first 10 stimulation trains and was increased every 10 stimulation trains by 50% of the initial current intensity. This strategy allowed to minimize the reduction of force production due to the repeated application of electrical pulses (Fig. 1E-F) and to maintain a mean force production (*i.e.*, training intensity) around 15% of F_max_ for each NMES session (Fig. 1G-H). The training intensity expressed in percentage of F_max_ slightly varied between mice (*i.e.*, ranging from ~7% to ~ 20% of F_max_; Fig. 1G-H) and between sessions (Fig. 1I-J & Table 1). The mean training intensity was 12.6 ± 2.7% of F_max_ and 13.6 ± 2.6% of F_max_ in C57BL/6J and BALB/c mice, respectively. The mean current intensity was 5.5 ± 1.4 mA and 5.5 ± 1.3 mA in C57BL/6J and BALB/c mice, respectively.

**TABLE 1.** Mean training intensity (expressed in percentage of maximal force; F_max_) and mean current intensity (expressed in mA) recorded during each NMES training session in C57BL/6J and BALB/c mice.

|  | Session 1 | Session 2 | Session 3 | Session 4 | Session 5 | Session 6 |
| --- | --- | --- | --- | --- | --- | --- |
| **C57BL/6J** |  |  |  |  |  |  |
| Training intensity (in % $F_{\max}$) | 12.5 ± 2.1 | 11.7 ± 1.9 | 12.3 ± 3.2 | 13.6 ± 1.9 | 11.7 ± 3.1 | 13.9 ± 3.2 |
| Current intensity (in mA) | 5.0 ± 1.6 | 5.3 ± 1.3<sup>#</sup> | 4.7 ± 1.6<sup>#</sup> | 6.0 ± 1.2 | 5.6 ± 1.5 | 6.2 ± 0.9 |
| **BALB/c** |  |  |  |  |  |  |
| Training intensity (in % $F_{\max}$) | 10.9 ± 1.1<sup>£</sup> | 13.7 ± 1.5 | 12.3 ± 1.2<sup>§</sup> | 13.8 ± 3.6 | 15.3 ± 2.0 | 15.7 ± 2.7 |
| Current intensity (in mA) | 4.3 ± 1.1<sup>§</sup> | 5.0 ± 1.2 | 4.8 ± 1.4 | 6.2 ± 1.1 | 6.3 ± 0.7 | 6.5 ± 0.6 |

NMES training was performed in n = 11 C57BL/6J mice and n = 6 BALB/c mice. <sup>#</sup>Significantly different from session 6: P< 0.05. <sup>§</sup>Significantly different from session 5: P< 0.05. <sup>£</sup>Significantly different from sessions 2; 3; 5 and 6: P< 0.05. Data were obtained from two to three separate experiments. Values are reported as mean ± SD.

### NMES training promotes myonuclear accretion and increased MuSC content

*Gastrocnemius* cryosections were immunostained for laminin and Hoechst in both NMES and control mice in order to evaluate the effects of the above described individualized NMES training sessions on myonuclear accretion. Nuclei with their geometric center within the inner rim of the laminin ring were defined as myonuclei (Fig. 2A-D; arrowheads). The number of myonuclei *per* fiber significantly increased by 21% and by 26% in C57BL/6J and BALB/c NMES trained mice as compared with controls, respectively (Fig. 2E-F). These results clearly demonstrate a robust myonuclear accretion after only 6 NMES training sessions performed at a submaximal force level of ~15% of F_max_.

**Figure 2.**
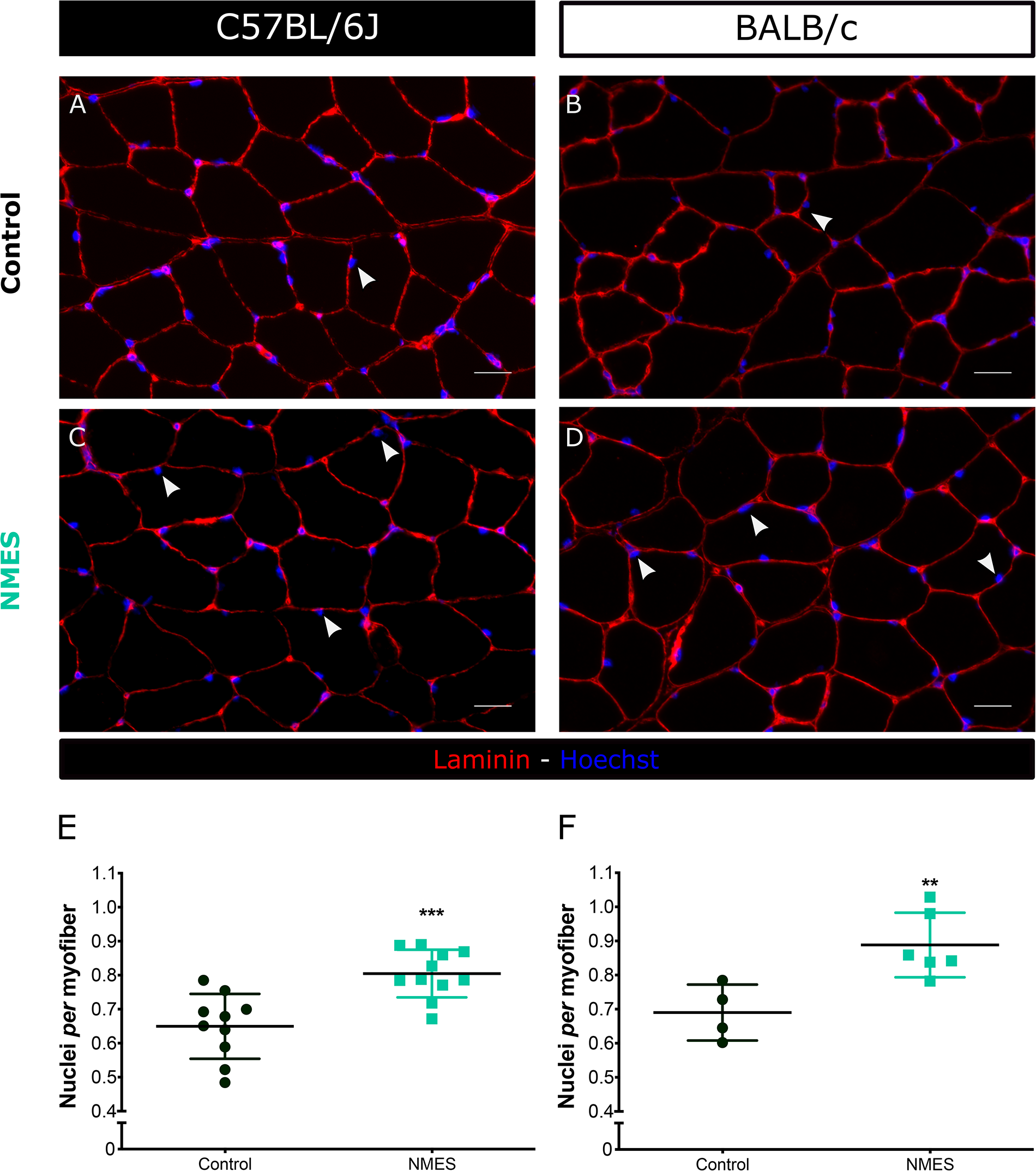
**A-D)** Immunostaining for laminin (red) and Hoechst (blue) on *gastrocnemius* muscle section from control and NMES trained mice. Arrowheads show myonuclei, *i.e.*, nuclei with their geometric center within the inner rim of the laminin ring. Scale bar = 25 μm. **E)** Number of myonuclei *per* myofiber in C57BL/6J control (n=10) and NMES trained (n=11) mice. **F)** Number of myonuclei *per* myofiber in BALB/c control (n=4) and NMES trained (n=6) mice. Significantly different from control: ^**^P< 0.01; ^***^P<0.001. Data were obtained from two to three separate experiments Values are reported as mean ± SD.

Next, we investigated whether NMES-induced myonuclear accretion was also associated with an increased number of MuSCs. *Gastrocnemius* cryosections were immunostained for Pax7 to label MuSCs in both NMES and control mice (Fig. 3A-D; arrowheads). The number of Pax7^+^ related to the number of myofibers increased by 67% (P<0.001) and by 48% (P=0.055) in C57BL/6J and BALB/c NMES trained mice as compared with controls, respectively (Fig. 3E-F), indicating that NMES increases the MuSC content in two different mouse lines.

**Figure 3.**
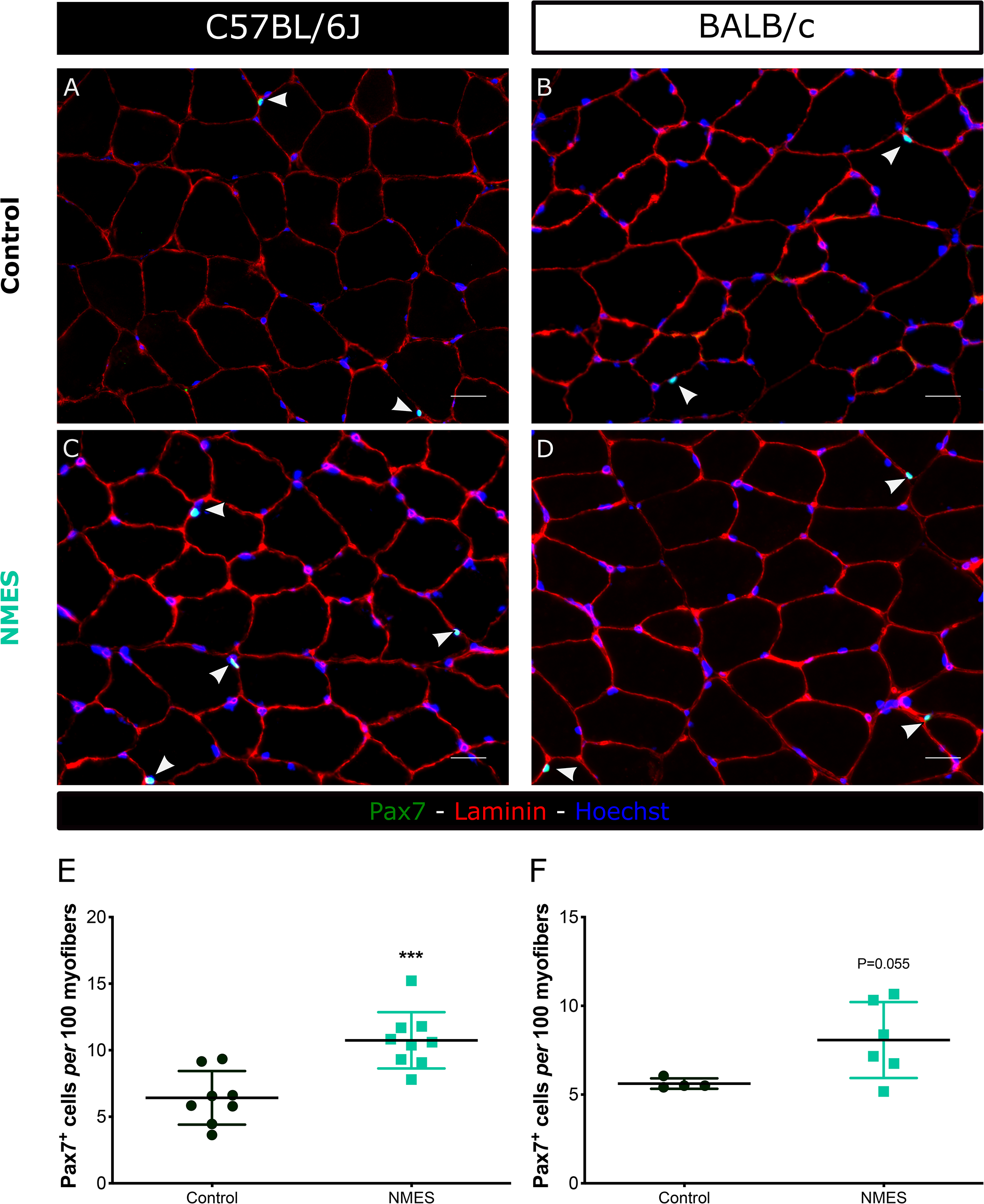
**A-D)** Immunostaining for laminin (red), Pax7 (green) and Hoechst (blue) on gastrocnemius muscle section from control and NMES trained mice. Arrowheads show nuclei positive for Pax7. Scale bar = 25 μm. **E)** Number of Pax7^+^ cells *per* 100 myofibers in C57BL/6J control (n=8) and NMES trained (n=9) mice **F)** Number of Pax7^+^ cells *per* 100 myofibers in BALB/c control (n=4) and NMES trained (n=6) mice. Significantly different from control: ^***^P<0.001. Data were obtained from two to three separate experiments. Values are reported as mean ± SD.

### NMES training does not induce overt signs of muscle damage

Considering that myonuclear accretion may be driven by muscle damage and/or regeneration (Murach *et al*, 2020a), we immunostained IgG trapping on *gastrocnemius* cryosections of both NMES and control mice as an index of increased membrane permeability (Supplemental Fig. 1). The proportion of myofibers positive for IgG was negligible in both NMES and control mice (*i.e.*, < 0.5%) (Fig. 4A-B; Supplemental Fig. 1A-B), suggesting that NMES does not induce membrane leakage.

**Figure 4.**
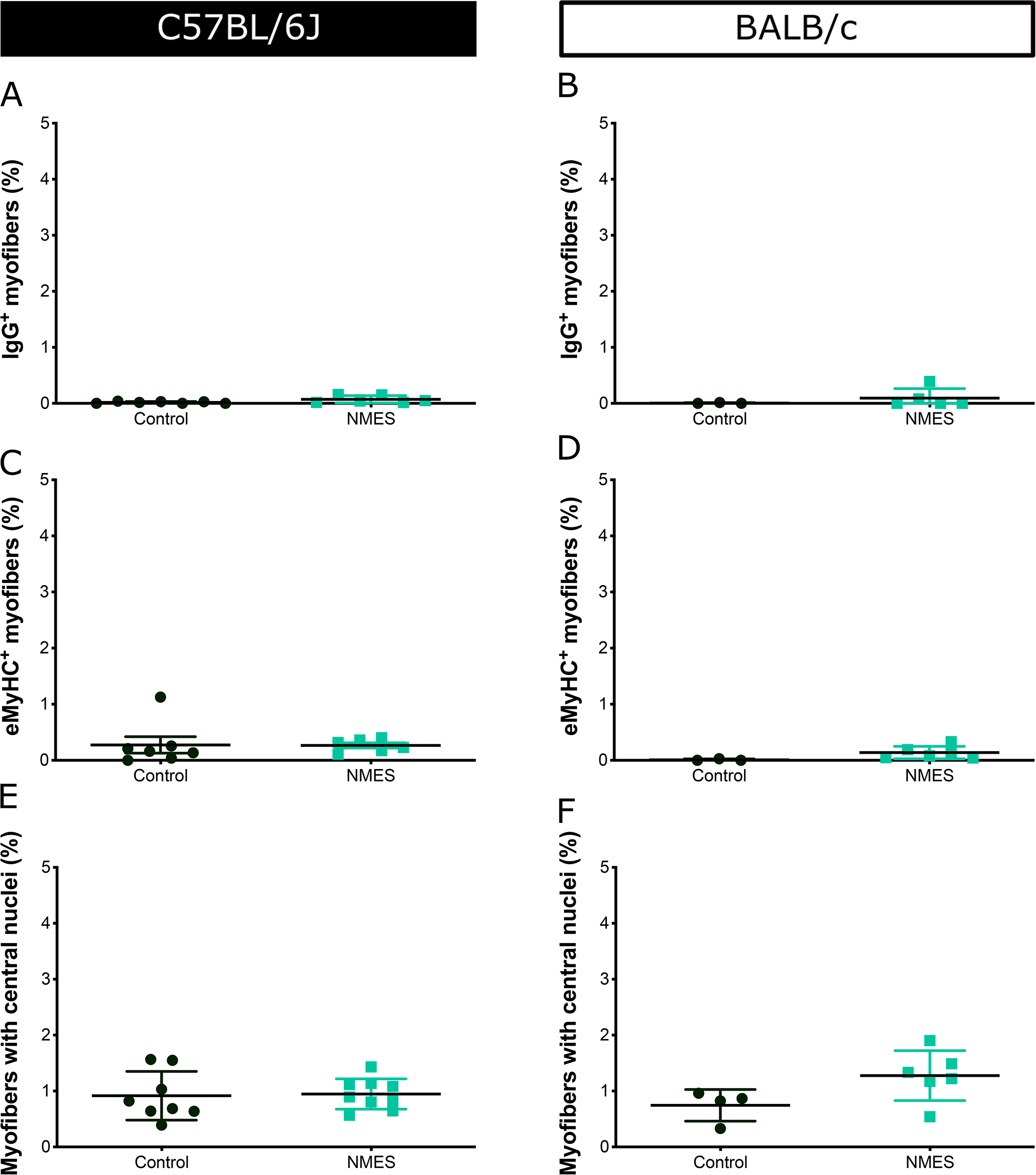
**A)** Proportion of myofibers positive for IgG in C57BL/6J control (n=7) and NMES trained (n=6) mice. **B)** Proportion of myofibers positive for IgG in BALB/c control (n=3) and NMES trained (n=5) mice. **C)** Proportion of myofibers positive for embryonic myosin heavy chain in C57BL/6J control (n=7) and NMES trained (n=6) mice. **D)** Proportion of myofibers positive for embryonic myosin heavy chain in BALB/c control (n=3) and NMES trained (n=6) mice. **E)** Proportion of myofibers with central nuclei in C57BL/6J control (n=8) and NMES trained (n=9) mice. **F)** Proportion of myofibers with central nuclei in control (n=4) and NMES trained (n=6) BALB/c mice. Data were obtained from two to three separate experiments. Values are reported as mean ± SD.

We also investigated whether signs of muscle regeneration can be observed in *gastrocnemius* muscles of both NMES and control mice. Cryosections were immunostained for embryonic MyHC (eMyHC) that labels newly formed myofibers, and laminin. The proportion of myofibers positive for eMyHC was counted on the whole section. The percentage of myofibers positive for eMyHC was very low (*i.e.*, < 1%) in both control and NMES mice and for the two different mouse lines (Fig. 4C-D; Supplemental Fig. 1C-D). Finally, we counted the number of myofibers with central nuclei, as central positioning of nuclei in myofibers is commonly used as a marker of regeneration. In agreement with the eMyHC staining, the percentage of myofibers with central nuclei was very low (*i.e.*, < 1-2%) in all mice (Fig. 4E-F). Overall, on the basis of those three different markers of muscle damage/regeneration, our results illustrate the non-damaging effects of NMES on *gastrocnemius* muscle.

### NMES does not induce muscle hypertrophy or force improvement

Considering that myonuclear accretion has been reported as an early event occurring before (or in absence of) muscle hypertrophy (Masschelein *et al*, 2020; Goh *et al*, 2019), *gastrocnemius* muscle weight and myofiber CSA were quantified in both control and NMES mice as indices of muscle hypertrophy. These two parameters were not significantly different between control and NMES mice in the two mouse lines, indicating that myonuclear accretion was not associated with an increase in muscle mass and myofiber size (Fig. 5A-D).

**Figure 5.**
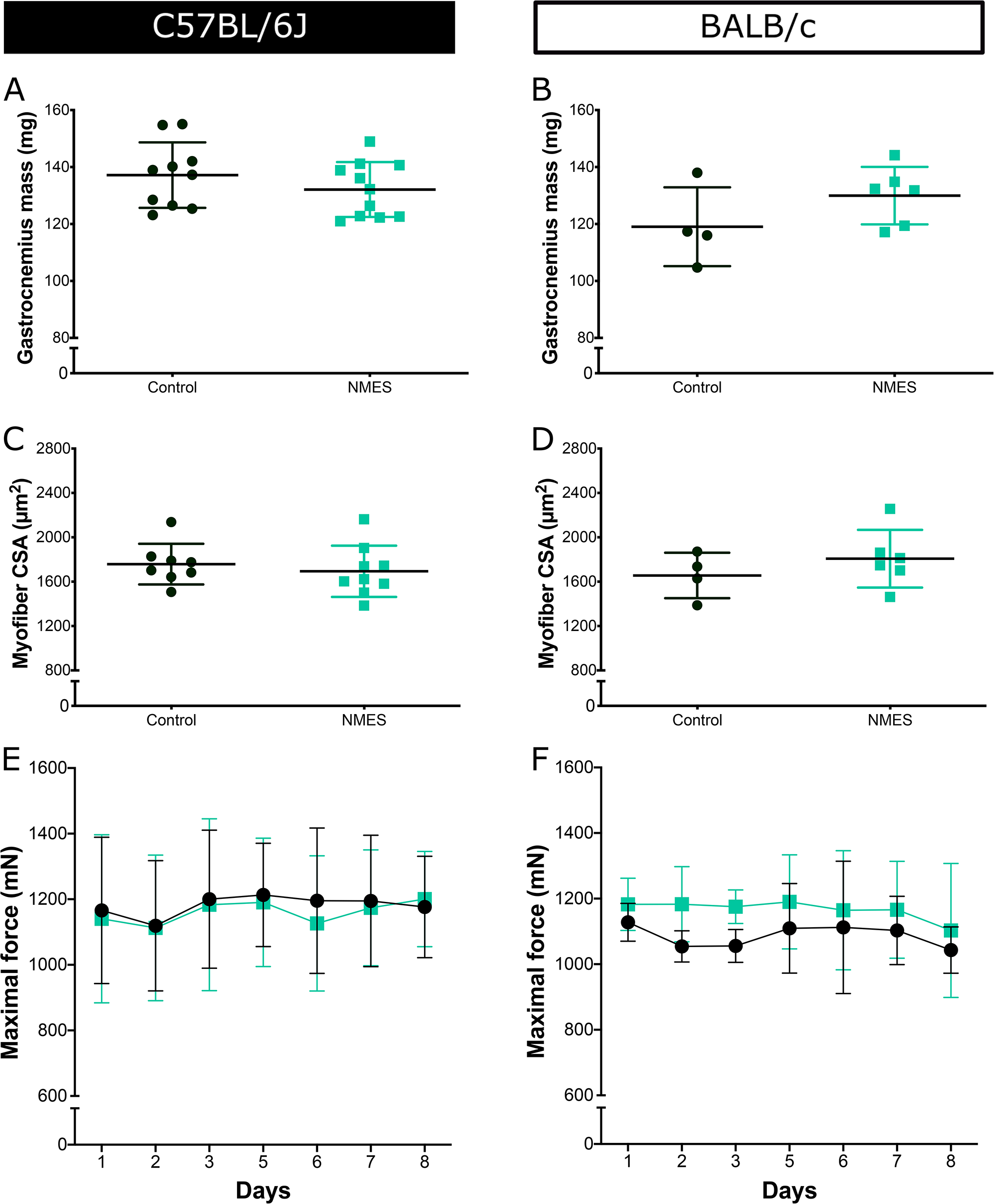
**A)** *Gastrocnemius* muscle weight of C57BL/6J control (n=10) and NMES trained (n=11) mice. **B)** *Gastrocnemius* muscle weight of BALB/c control (n=4) and NMES trained (n=6) mice. **C)** *Gastrocnemius* myofiber cross-sectional area of C57BL/6J control (n=8) and NMES trained (n=9) mice. **D)** *Gastrocnemius* myofiber cross-sectional area of BALB/c control (n=4) and NMES trained (n=6) mice. **E)** Maximal isometric force production longitudinally recorded throughout the study design in C57BL/6J control (n=10; black circles) and NMES trained (n=11; green squares) mice. **F)** Maximal isometric force production longitudinally recorded throughout the study design in control (n=4; black circles) and NMES trained (n=6; green squares) BALB/c mice. Data were obtained from two to three separate experiments. Values are reported as mean ± SD.

Finally, we investigated whether the changes in both myonuclear and MuSC content might have a functional effect in terms of muscle force production. Maximal tetanic force (*i.e.*, F_max_) was recorded longitudinally in both control and NMES mice throughout the duration of the experiment (*i.e.*, Fig. 1B). Our functional analysis reveals that six NMES training sessions did not increase muscle force in either C57BL/6J and BALB/c mice (Fig. 5E-F).

## DISCUSSION

In the present study, we took advantage of our original device allowing for non-invasive force measurements in response to electrical stimuli applied over the plantar flexor muscle belly to design individualized and carefully monitored isometric NMES training sessions in two mouse lines. We observed an elevation of both nuclei *per* myofiber and MuSC content after only six isometric NMES training sessions performed at submaximal force level of 15% of F_max_. These changes were not associated with overt signs of muscle damage/regeneration, and were not accompanied by muscle hypertrophy or force improvement Thereby, we demonstrate that myonuclear accretion is a rapid process occurring prior to, or in the absence of muscle hypertrophy and force improvement.

The individualized isometric NMES training program led to a robust myonuclear accretion as illustrated by the 21% and 26% increase in number of nuclei *per* myofiber in C57BL/6J and BALB/c mice, respectively. These results are consistent with the ~15-25% increase in myonuclei *per* fiber reported in different muscles (e.g., *soleus, gastrocnemius, plantaris*) after voluntary wheel running (Dungan *et al*, 2019; Masschelein *et al*, 2020) or high-intensity interval treadmill (Goh *et al*, 2019). However, it is worth noting that the magnitude of myonuclear accretion is higher after NMES than after running exercise. Indeed, the duration of NMES-induced contractile activity was less than 3 min *per* session (*i.e.*, 80 stimulation trains lasting 2 s) for a total contractile activity of ~16 min over a one-week training period while treadmill or wheel running exercises usually involve 60-300 min session duration with 3-7 sessions per week for ~8 weeks (Murach *et al*, 2020a; Goh *et al*, 2019; Masschelein *et al*, 2020). One can therefore estimate that the increase of myonuclei per fiber per min of stimulation/exercise is ~100-fold higher after NMES as compared with running exercise. Interestingly, NMES-induced myonuclear accretion was associated with a large increase in the number of MuSCs present in the trained muscle (*i.e.*, +~ 50-70%). This is also consistent with the results obtained after voluntary wheel running (+44% (Dungan *et al*, 2019)), even though changes in MuSC content are also more pronounced after NMES as compared with running protocols. Our innovative NMES training protocol appears therefore as a method of choice for investigating the contribution of MuSCs to myonuclear accretion. Moreover, the use of genetically engineered mouse models ablated for MuSCs (Egner *et al*, 2016; McCarthy *et al*, 2011), deleted for *myomaker* (Goh & Millay, 2017) or allowing the labeling of MuSCs (Pawlikowski *et al*, 2015; Keefe *et al*, 2015; Masschelein *et al*, 2020) would allow to decipher fusion -dependent and/or -independent MuSC communication with myofibers (Murach *et al*, 2020b) in response to NMES.

The present study also demonstrates that NMES-induced myonuclear accretion occurs in the absence of overt signs of muscle damage/regeneration. Indeed, the proportion of myofibers positive either for IgG labeling or embryonic MyHC as well as the percentage of myofibers with central nuclei was very low (*i.e.*, < 1-2%) and was never different between control and NMES mice, independent of their genetic background. On the contrary, the proportion of myofibers with central nuclei was significantly higher in trained mice as compared with control animals in response to wheel running exercise (Murach *et al*, 2020a; Masschelein *et al*, 2020) and reached a mean value of ~6% (range: ~2-10% in the soleus muscle; (Murach *et al*, 2020a)) with (Murach *et al*, 2020a) or without (Masschelein *et al*, 2020) the co-expression of embryonic MyHC. This indicates that the individualized isometric NMES training protocol performed at low force levels (*i.e.*, ~10-15% of F_max_) can be considered as a non-damaging modality of increased mechanical loading, whereas running exercise contains a component of damaging eccentric contractions. This is further supported by our functional analyses showing that NMES does not alter maximal force production, this parameter being considered as the best indirect marker of muscle damage (Warren *et al*, 1999). Our data therefore suggest that myonucler accretion is primarily mediated by NMES-induced myofiber contractile activity rather than muscle damage. Further studies are warranted to identify factors secreted by MuSCs (Fry *et al*, 2017) and/or contracting myofibers (Guerci *et al*, 2012) involved in NMES-induced skeletal muscle remodeling.

Our results also show that NMES-induced myonuclear accretion and higher MuSC content did not increase *gastrocnemius* muscle weight and myofiber size. This lack of muscle hypertrophy can be explained by the short training duration (*i.e.*, only 6 NMES sessions over one week corresponding to a total contractile activity of ~16 min) and/or the submaximal training intensity (*i.e.*, only ~10-15% of F_max_). Indeed, we previously reported an increase in human quadriceps muscle and myofiber size after 8 weeks of NMES performed at 55-60% of F_max_ (Gondin *et al*, 2005, 2011b). Additional studies are required to investigate whether longer NMES training duration and/or higher training intensity could induce hypertrophy in mice. Our data further demonstrate that NMES-induced myonuclear accretion is an early event occurring in the absence of myofiber CSA changes. This is in agreement with previous studies showing that myonuclear accretion precedes muscle hypertrophy in response to mechanical overload (Bruusgaard *et al*, 2010) or treadmill exercise (Goh & Millay, 2017).

In the present study, experiments were performed in mice with two different genetic backgrounds (*i.e.*, C57BL/6J and BALB/c) that are commonly used to investigate the impact of cancer cachexia (thanks to the inoculation of LLC or C26 tumor cells; (Gallot *et al*, 2014; Penna *et al*, 2016)) or sepsis (*i.e.*, using cecal ligation and puncture (Morel *et al*, 2017)) on skeletal muscle homeostasis. Indeed, recent evidence is emerging regarding the key role of defective regulation of MuSCs on cancer cachexia-(He *et al*, 2013) or sepsis-induced (Rocheteau *et al*, 2015) muscle atrophy. On that basis, NMES could be a relevant method to promote MuSC fusion in these two pathological contexts. However, additional investigations are warranted to determine whether and to what extent NMES is a relevant non-pharmacological approach to minimize the deleterious consequences of cancer cachexia or sepsis on the regulation of MuSCs.

In conclusion, we provide here an individualized and carefully-controlled isometric NMES training protocol that allows to promote a robust myonuclear accretion and an increase in MuSC content without inducing muscle damage/regeneration or hypertrophy. We demonstrate that myonuclear accretion is primarily driven by increased mechanical loading rather than muscle injury. This new mouse model of myonuclear accretion that relies on the main function of skeletal muscle, *i.e.*, force production in response to electrical stimuli, will be of utmost interest to further understand the role of MuSCs in skeletal muscle adaptations.

## Author Contributions

Concept/idea/research design: J. Gondin

Writing: J. Gondin

Data collection: A. Zavoriti, A. Fessard, M. Rahmati, P. Del Carmine, J. Gondin

Data analysis: A. Zavoriti, A. Fessard, P. Del Carmine, J. Gondin

Project management: J. Gondin

Consultation (including review of manuscript before submitting): A. Zavoriti, A. Fessard, M. Rahmati, P. Del Carmine, B. Chazaud, J. Gondin

All the authors read and approved the final version of manuscript. Masoud Rahmati acted as a visiting scientist at INMG (from September 2019 to July 2020) where he was trained to NMES training protocol and muscle analysis.

## Acknowledgments

We thank Dr Rémi Mounier and Dr Anita Kneppers for their useful comments on a previous version of the manuscript.

## Supplemental Materials

**Supplemental video 1.** A video clip illustrating the positioning of a BALC/c mouse in the ergometer. The application of electrode cream on the right plantar flexor muscles as well as the position of the foot on the pedal are also shown.

**Supplemental Figure 1. A-B)** Immunostaining for IgG (red) and laminin (green) on gastrocnemius muscle section from C57BL/6J control and NMES trained mice. Scale bar = 250 μm. **C-D)** Immunostaining for laminin (red) and embryonic myosin heavy chain (green) on *gastrocnemius* muscle section from C57BL/6J control and NMES trained mice. Scale bar = 250 μm.

