## Supplementary figures and images for "Individualized isometric neuromuscular electrical stimulation training promotes myonuclear accretion in mouse skeletal muscle"

### Supplemental figure 1

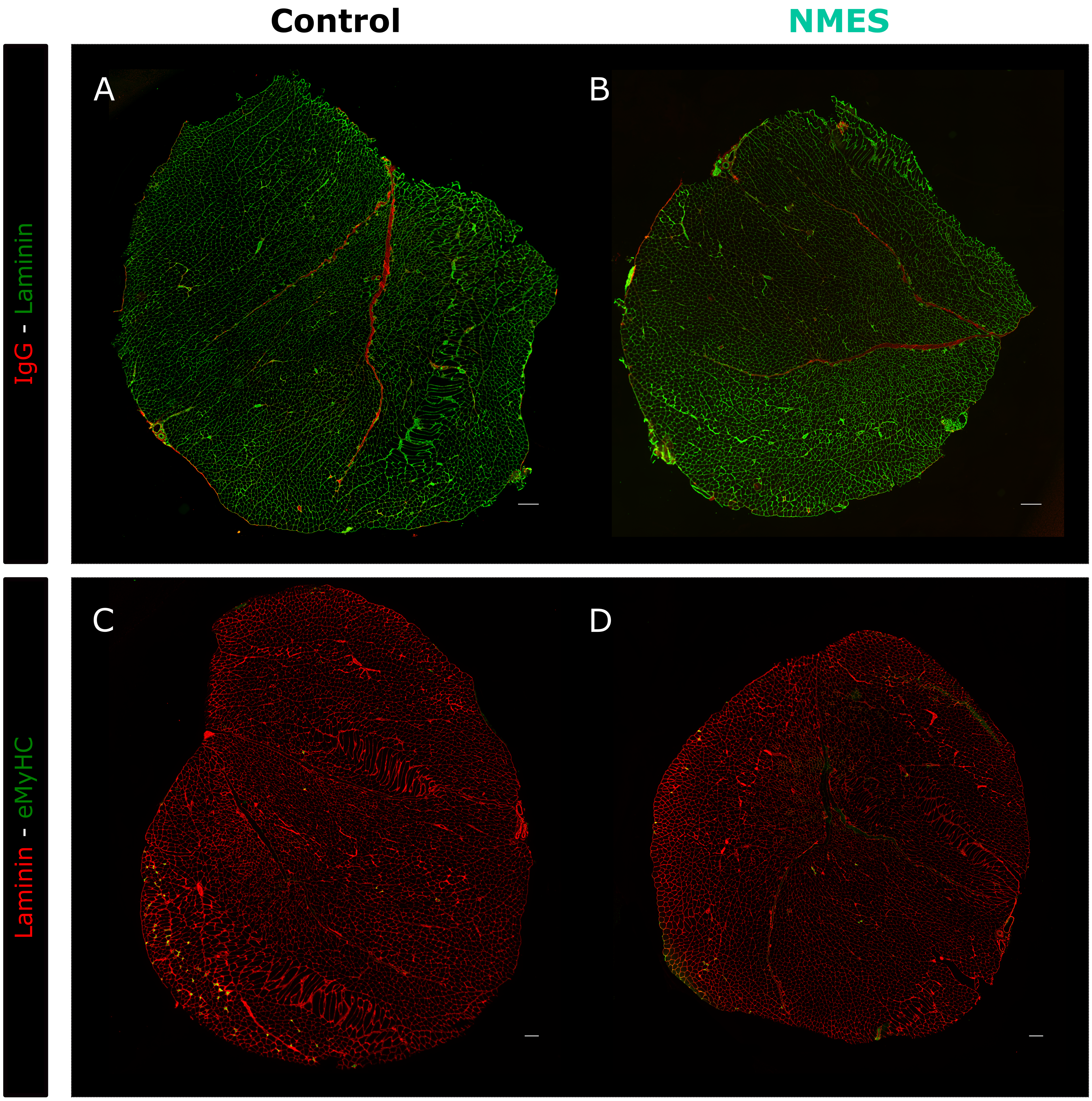
